# The missing pathway in current visuospatial processing models

**DOI:** 10.1101/2021.06.08.442147

**Authors:** Ahmad Beyh, Flavio Dell’Acqua, Daniele Cancemi, Francisco De Santiago Requejo, Dominic ffytche, Marco Catani

**Author notes:** Corresponding author: Ahmad Beyh, PO50 Institute of Psychiatry, Psychology and Neuroscience, King’s College London, De Crespigny Park, London SE5 8AF, UK.

## Abstract

Visuospatial learning depends on the parahippocampal place area (PPA), a functionally heterogenous area which current visuospatial processing models place downstream from parietal cortex and only from area V4 of early visual cortex (EVC). However, evidence for anatomical connections between the PPA and other EVC areas is inconsistent, and these connections are not discussed in current models. Through a data-driven analysis based on diffusion MRI tractography, we present evidence that the PPA sits at the confluence of two white matter systems. The first conveys information from the retrosplenial complex to the anterior PPA and runs within the cingulum bundle. The second system connects all peripheral EVC areas to the posterior PPA and corresponds to the medial occipital longitudinal tract (MOLT), a novel white matter pathway distinct from the cingulum. Based on further functional connectivity analysis and meta-analytic data, we propose that the MOLT supports early stage encoding of visuospatial information by allowing direct reciprocal exchange between the PPA and EVC.

## INTRODUCTION

Multiple streams of research into rodents, non-human, and human primates have identified several brain regions that contribute to the complex ability of visuospatial learning. Although the hippocampal formation is a key player in this cognitive domain ^1–3^, several reports suggest that the more posterior parahippocampal gyrus (PHG) lodges cortical areas more vital for specific types of visuospatial learning, including configuration learning ^4–8^. Configuration learning requires building associations between objects and their location in the visual space. As such, it has been partially dissociated from the memory for object identities and separate anatomical substrates have been identified for each in human ^7,9,10^ and non-human primates ^11^. Focal lesion studies have linked configuration learning to an area of the posterior collateral sulcus (CoS) known as the parahippocampal place area (PPA) ^7–9,12^. Conversely, memory for objects has been associated with the hippocampus and with areas distributed along the ventral visual pathway connecting occipital to ventrolateral temporal cortex ^13–15^.

Functional imaging investigations have revealed that responses within the PHG are organised along an anterior-to-posterior gradient, with anterior PHG being more involved in identity encoding, and responses to configuration changes occupying more posterior locations ^10^. Although early descriptions of the PPA focused on its preferential responses to visual stimuli depicting scenes ^12,16^ and treated it as a single functional unit, subsequent resting state fMRI connectivity analyses have revealed a similar gradient within the PPA itself based on its coupling to early visual cortex (EVC), parietal, and frontal regions ^17,18^. To date, it is unclear whether this functional gradient within the PPA is driven by local, intrinsic organisation or by external anatomical input.

A counterpart to the PPA that also plays a central role in the spatial learning domain is the retrosplenial complex (RSC). Although the retrosplenial cortex is an anatomical label which corresponds to Brodmann areas (BA) 29 and 30, the term RSC is used here to refer to the cortical region which overlaps the anatomical retrosplenial cortex and posterior cingulate cortex (PCC, BA 23 and 31) ^19^. Located immediately caudal to the splenium of the corpus callosum, the RSC is strategically located to receive body-centred (egocentric) spatial information from posterior parietal cortex (PPC) ^20,21^ and allocentric spatial information from regions of the anterior medial temporal lobe (MTL) ^22^. Unlike the PPA, the RSC responds much more to images of familiar scenes compared to novel scenes and its responses are viewpoint-independent ^23^, which supports the idea that the RSC is not performing simple spatial mapping but is rather involved in a more complex, memory-driven task ^24^.

Based on this and other evidence of anatomical connectivity, Kravitz et al. ^21^ revised the influential dual-stream model for visuo-spatial processing ^14^ to include, within the dorsal stream for spatial processing, a branch connecting the posterior parietal lobe to the MTL, including the hippocampus and PPA (see Rossetti et al. ^25^ for a criticism of the model). This pathway linking parietal and medial temporal cortices is known as the third branch of the dorsal stream, and projects both directly and indirectly to the MTL. While anterior MTL regions such as the hippocampal formation mainly receive direct parietal projections, input to the posterior PHG stems mainly from indirect connections relayed via the RSC and other regions of medial parietal cortex (MPC) (see Kravitz et al. ^21^ for a full review of these connections).

Deep brain electrode recordings in surgical patients have revealed responses to visual stimulation in the PPA as early as 80 ms ^26,27^. This suggests that EVC may be sending visual information directly to the PPA, bypassing the intricate polysynaptic circuitry of the dorsal visual stream. Complementary evidence from the axonal tracing literature supports this notion in the macaque. First, there is support for the existence of direct connections between peripheral visual representations in area V4 and the posterior PHG ^28–30^. Further, Gattas ^31^ injected anterograde tracers in the macaque ventral V2 and observed cortical projections in the most posterior portion of the PHG. Starting at the other end, Schmahmann and Pandya ^32^ reported V2 projections following anterograde tracer injections in the macaque posterior PHG. However, they did not label these fibres separately, possibly because they may be difficult to discern from those of the inferior longitudinal fasciculus in the macaque brain.

Recently, Catani used diffusion MRI tractography to report a ‘Sledge Runner’ fasciculus ^33^, and two human *post mortem* dissection reports successfully dissected the reported fasciculus ^34,35^. However, all these reports on humans described a white matter bundle that runs between the posterior PHG and the medial occipital wall, terminating in the most dorsal part of the cuneus. This disagrees with the macaque literature which mainly reports connections with ventral EVC.

Here, we address the inconsistent findings regarding these connections in the human brain using tractography based on diffusion MRI data from 200 participants from the Human Connectome Project (HCP, https://www.humanconnectome.org/). First, we first demonstrate that the pattern of the MTL’s anatomical connections to EVC and RSC/MPC is central to understanding its functional heterogeneity. Through a data-driven clustering analysis of structural connectivity tractography data, we demonstrate that the posterior MTL, specifically the PPA, sits at the confluence of two white matter systems: a dorso-medial system stemming from the RSC/MPC and a ventromedial system stemming from EVC. Anatomically, both systems merge with the posterior cingulum bundle and this may have prevented their identification in early tractography studies. Here were propose the separation of the ventral EVC-PPA system from the cingulum, and its identification as a novel pathway that we refer to as the medial occipital longitudinal tract (MOLT).

## RESULTS

### The PPA sits at the confluence of two white matter systems

As a first step, we sought to study the general connectivity pattern between the MTL and two sets of regions. The first set, based on the revised dual-stream model ^21,23,36,37^, included the RSC and additional surrounding regions from the MPC (RSC/MPC region in Figure 1). The second set of regions was defined to fully cover EVC within the occipital lobe (EVC region in Figure 1).

**Figure 1.**
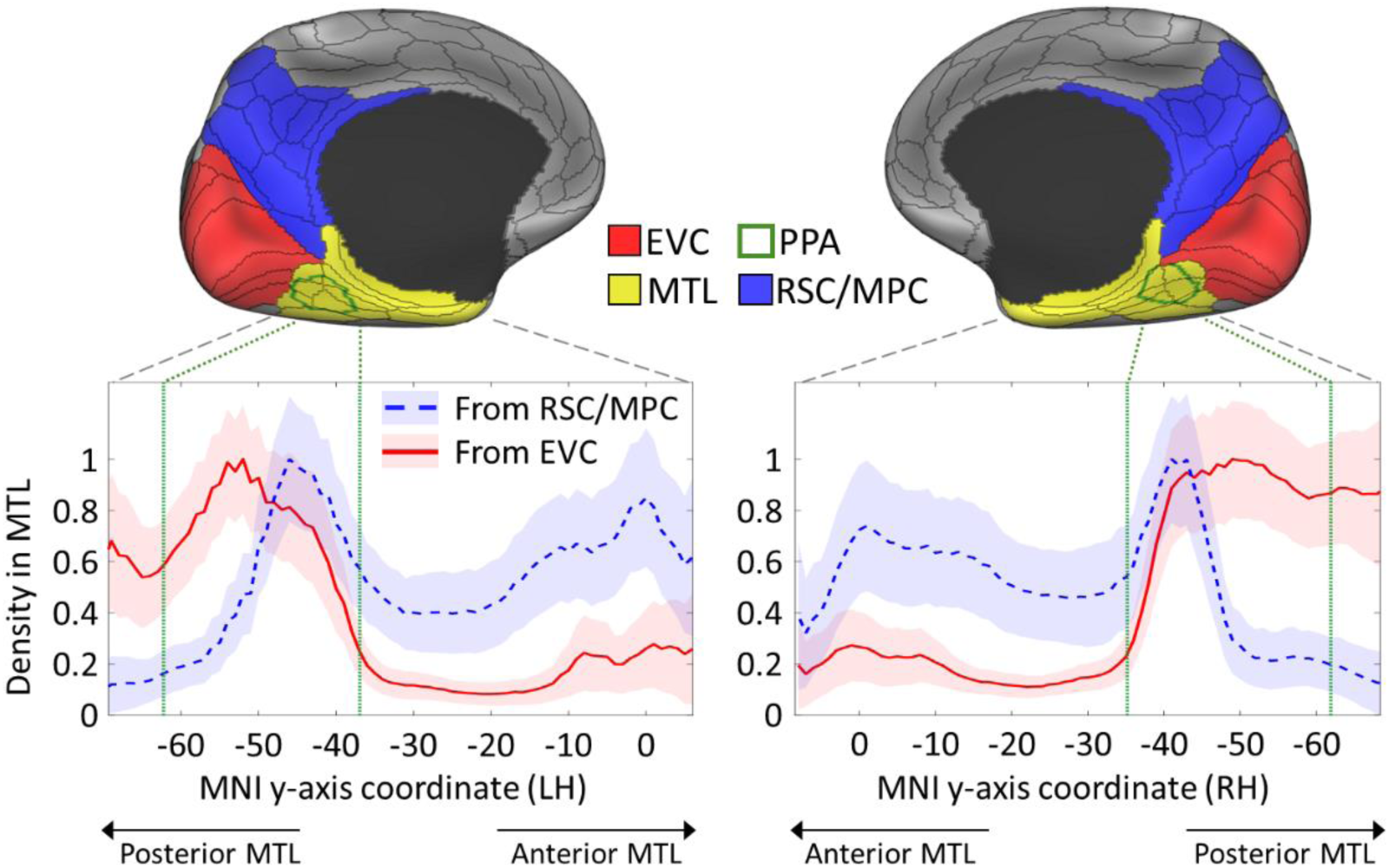
Density of tractography projections from medial occipital and parietal regions to the medial temporal lobe. The upper panel shows the three large ROIs used to select streamlines between the medial temporal lobe (MTL) on one end, and early visual cortex (EVC), medial parietal cortex (MPC) and retrosplenial complex (RSC) on the other end. The parahippocampal place area (PPA), which is located in the posterior MTL, is also delineated with a green contour. The plots in the lower panel represent the mean density of streamline terminations in the left and right MTL according to its longitudinal y-axis. Projections from both EVC (red lines) or the RSC/MPC (blue dashed lines) are displayed as the mean and standard deviation (shaded areas) across 200 subjects. While the EVC connections show a strong preference for the posterior MTL, especially the parahippocampal place area (PPA), the RSC/MPC connections show posterior and anterior peaks. Within the PPA, the anterior portion receives high density projections from both ROIs, while the posterior portion receives projection preferentially from EVC. The PPA’s most posterior and anterior coordinates are indicated with the vertical green lines within the plots. Coordinates correspond to the MNI template. The main ROIs are based on the MMP atlas shown as black contours ^38^, and the PPA is based on ^39^.

To select the tractography streamlines that connect these pairs of regions, we converted the whole-brain tractogram of each participant to a surface-based vertex-wise connectivity matrix. We then filtered these matrices to only keep connections between the MTL on one end, and either the EVC or RSC/MPC regions on the other end. This produced a vertex-to-vertex connectivity map for each pair of regions in each participant.

Figure 1 shows the mean density of the observed connections in the MTL in each hemisphere. On one hand, there is a clear trend where the posterior MTL receives the majority of EVC connections, followed by a sharp decrease moving toward the anterior MTL. This trend is statistically significant according to a Spearman rank correlation: left hemisphere, *r*_*S*_ = -.72, *p* < .001; right hemisphere *r*_*S*_ = -.67, *p* < .001. On the other hand, the RSC/MPC connections show prominent peaks in both the posterior and anterior MTL, resulting in a non-significant trend overall: left hemisphere, *r*_*S*_ = .35, *p* = 1.00 (*n*.*s*.); right hemisphere *r*_*S*_ = .39, *p* = 1.00 (*n*.*s*.).

In both cases, the zone with the highest density coincides with the location of the PPA but the density of these connections within the PPA is not homogeneous. For the EVC connections, the highest density is observed in the posterior half of the PPA, with a decline to near-zero density towards its anterior border (Figure 1). However, the density of RSC/MPC connections is negligible in the posterior half of the PPA and sharply rises to its highest peak in the anterior PPA. These findings suggest that the functional gradients that exist within the PPA ^17,18^ are driven by an interaction between two anatomical systems stemming from EVC and RSC/MPC.

### Clustering analysis of the PPA’s connections

To better understand how the two observed white matter systems interact structurally and functionally within the PPA, we resorted to a data-driven clustering analysis. To this end, we first combined the EVC and RSC/MPC regions into a single large ROI, and filtered the structural connectivity matrices to only keep connections between the PPA on one end and this combined ROI on the other end. Next, we used principal component analysis (PCA) to only retain the main features that represent these connections. In both hemispheres, PCA yielded three principal components which explained 89.1% (LH) and 89.8% (RH) of the variance in the connectivity profile of the PPA to the target ROI (Figure 2).

**Figure 2.**
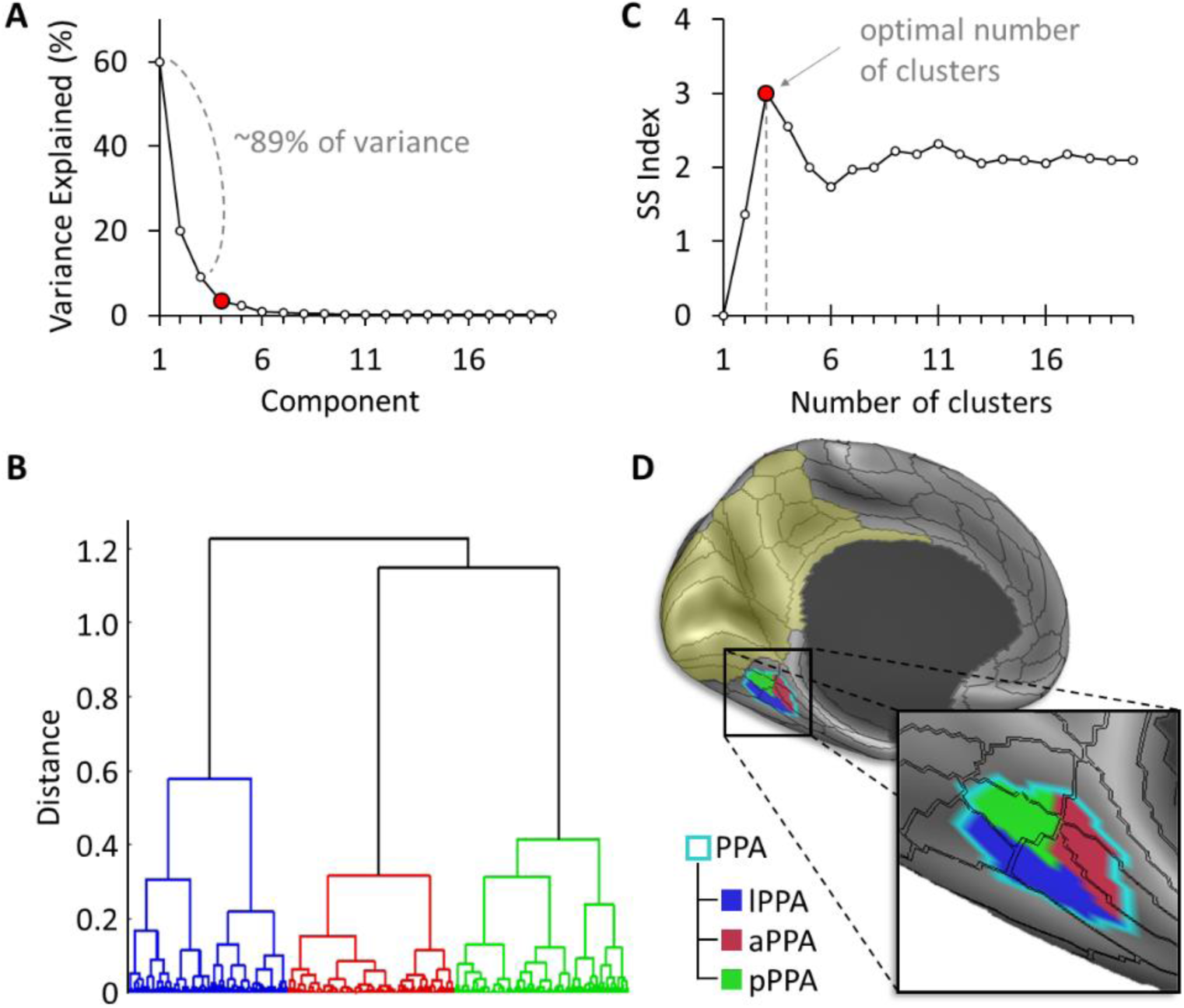
Clustering analysis of the PPA reveals multiple anatomical subunits. Data-driven clustering of the parahippocampal place area (PPA) based on structural connectivity. **(A)** Principal component analysis (PCA) based on the PPA’s connectivity to a region encompassing early visual cortex (EVC), the retrosplenial complex (RSC), and medial parietal cortex (MPC) (yellow tint) resulted in three principal components. **(B)** Hierarchical agglomerative clustering grouped PPA surface vertices with similar PCA coefficients. **(C)** The highest separation vs. spread (SS) index ^40^ objectively determined the optimal number of clusters. **(D)** The resulting anterior, posterior, and lateral PPA clusters are shown on the inflated brain surface.

We then applied a hierarchical clustering analysis to extract distinct connectivity-based clusters within the PPA based on the identified principal components (Figure 2). We used the separation vs. spread (SS) index ^40^ to determine the clustering’s optimal granularity. In both hemispheres, a granularity of three clusters yielded the highest SS index: *SS*_*max*_ = 2.71 (LH); *SS*_*max*_ = 3.01 (RH). This indicates that the PPA is best subdivided into three clusters based on its structural connectivity to the region encompassing EVC, RSC, and the MPC.

In Figure 3, the structural connectivity of the three PPA clusters to EVC and RSC/MPC is shown and compared to their functional connectivity, which is based on an average resting-state fMRI connectivity map obtained from an HCP dataset of 812 subjects (analysed and released by the HCP; see Methods section). On one hand, the anterior cluster (aPPA) is mainly connected to the RSC/MPC according to both structural and functional connectivity. A Spearman rank correlation confirmed the agreement between the two methods: *r*_*s*_ = .57, *p* < .001 (LH); *r*_*s*_ = .72, *p* < .001 (RH). On the other hand, the posterior cluster (pPPA) is mainly connected to peripheral visual representations in EVC areas V1, V2 and V3, with a strong agreement between structural and functional connectivity: *r*_*s*_ = .61, *p* < .001 (LH); *r*_*s*_ = .65, *p* < .001 (RH).

**Figure 3.**
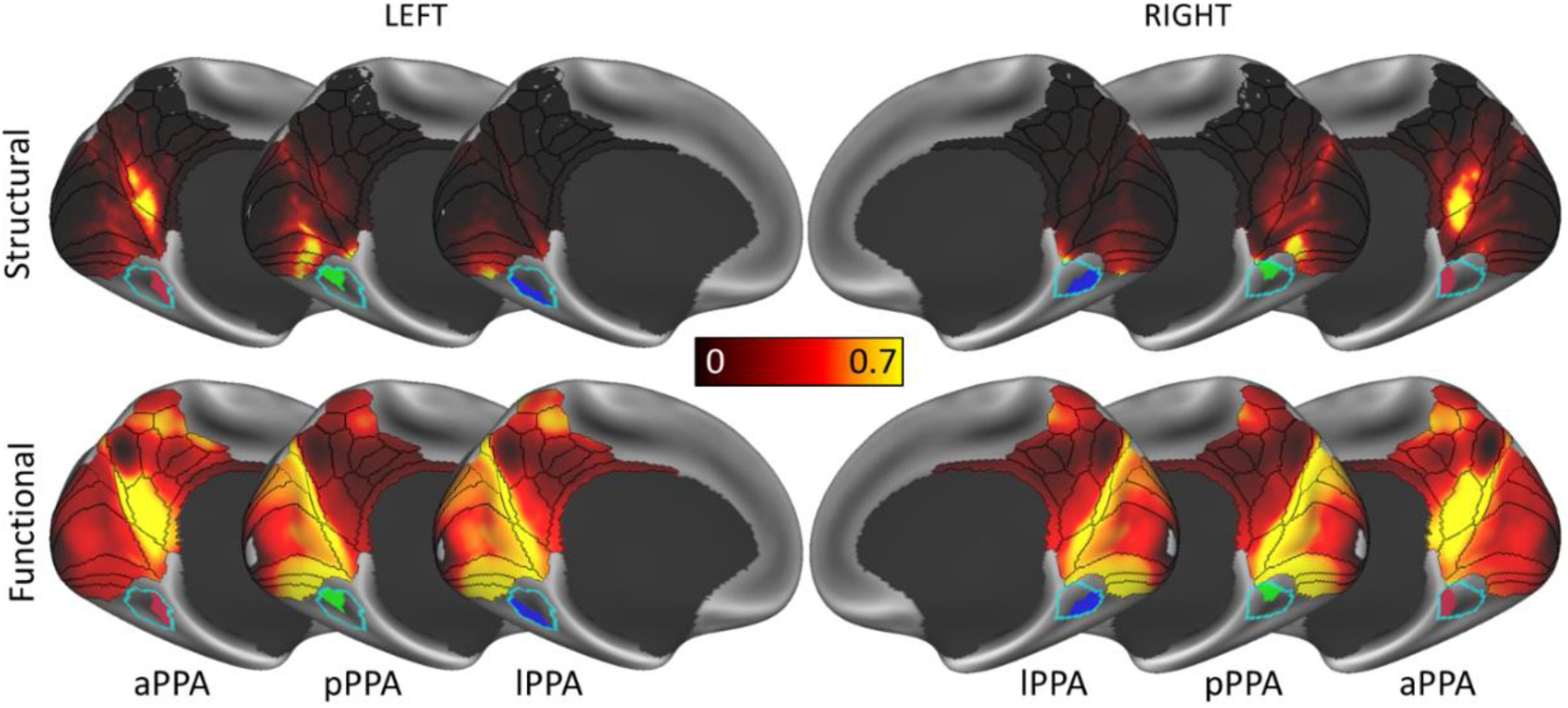
PPA clusters share similar structural and functional connectivity profiles. Structural (top row) and functional (bottom row) connectivity of the three parahippocampal place area (PPA) clusters. The anterior cluster (aPPA) is preferentially connected to retrosplenial complex and corresponds to the parieto-medial-temporal branch of the dorsal visual stream ^21^. The posterior cluster (pPPA) is preferentially connected to the anterior medial occipital lobe, i.e., peripheral representations within early visual cortex (EVC). The lateral cluster (lPPA) is more ambiguous but is preferentially connected to EVC (this is discussed in the Supplemental Material).

Although there is strong agreement between structural and functional connectivity for the anterior and posterior clusters, with a clear distinction between the two clusters, the correspondence between structural and functional connectivity of the lateral cluster (lPPA) is more ambiguous. While it shows preferential structural connectivity to area V4 of EVC, its functional connectivity appears more similar to that of the pPPA. This ambiguity is captured by the weakest agreement between structural and functional connectivity for this cluster: *r*_*s*_ = .45, *p* < .001 (LH); *r*_*s*_ = .51, *p* < .001 (RH). An in-depth discussion of this finding and further statistical comparisons are available in the Supplemental Material.

### From connectivity to functional specialisation

The results of the structural and functional connectivity analyses indicate that the RSC/MPC and EVC anatomical systems feed into the anterior and posterior PPA, respectively. This suggests that the functional specialisation within the PPA should reflect the distinct roles of RSC/MPC and EVC in the encoding and retrieval of visuospatial information ^41^. To address this point, we resorted to a meta-analysis ^42^ of PPA activations for the terms ‘encoding’ and ‘retrieval’ obtained from NeuroSynth (https://neurosynth.org/).

Figure 4 shows the distribution of the maximum z-score attributed to each of the terms ‘encoding’ and ‘retrieval’ along the posterior-anterior axis of the PPA. In the left hemisphere, encoding and retrieval show a similar trend: both have a low load posteriorly (z-score < 2) and an increasingly higher load moving anteriorly. However, while both encoding and retrieval are associated with the anterior PPA in the right hemisphere, only encoding shows a high peak in the posterior PPA (z-score > 2 starting at y = -60 mm) and its overall load in the right PPA is much higher. These results are in line with previous functional imaging studies and meta-analyses ^24,43^.

**Figure 4.**
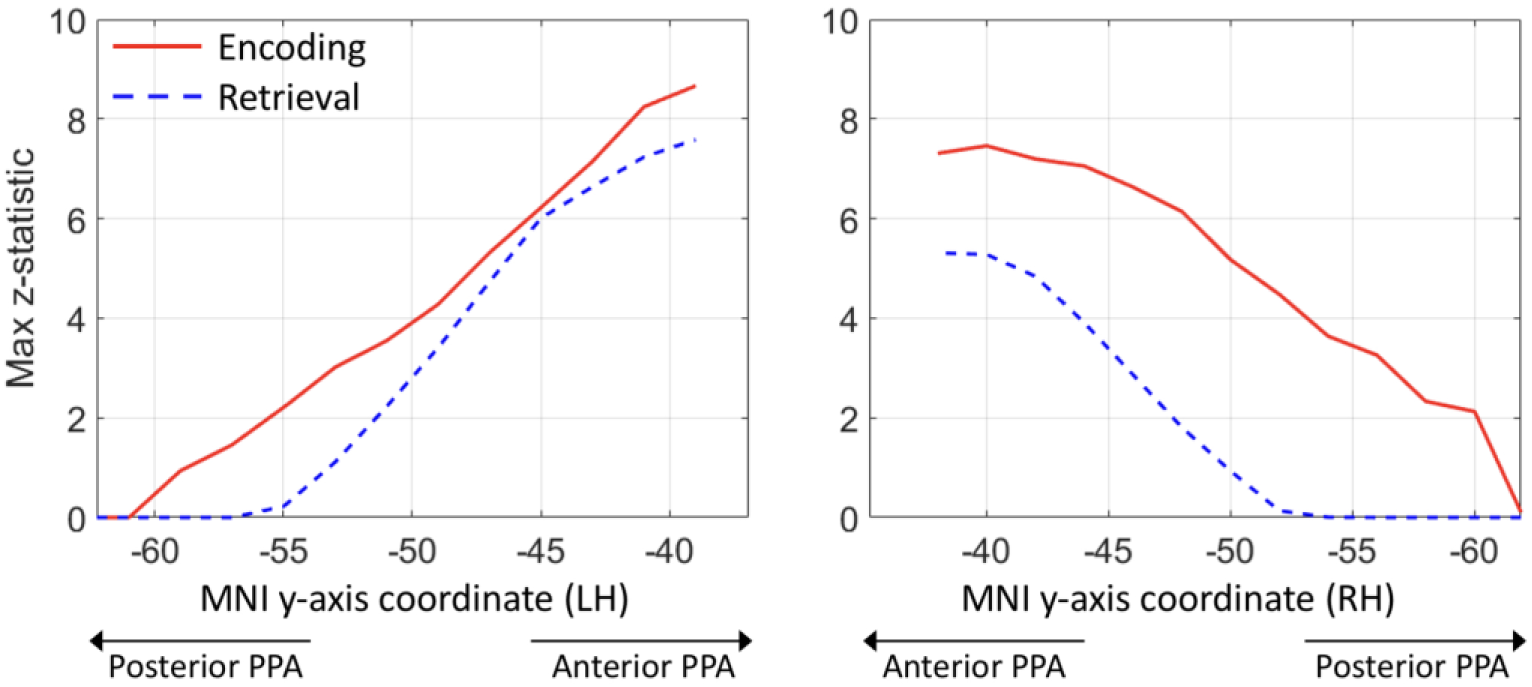
Encoding and retrieval in the PPA. These plots are based on meta-analytic maps, obtained from NeuroSynth, of locations associated with the terms ‘encoding’ and ‘retrieval’. The maximum z-statistic is plotted along the posterior-anterior axis of the parahippocampal place area (PPA).

### White matter bundle connecting the PPA to EVC

The connectivity analyses and the meta-analytic results point to the existence of at least two anatomical systems converging within the PPA. While the PPA’s connections to the RSC/MPC have been well characterised and described within the posterior cingulum bundle and are well discussed in current models, the anatomy of the connections between the PPA and EVC is less understood. On that account, we performed semi-automatic tractography dissections ^44^ of the latter connections in the 200 HCP datasets using the ROIs described in the Methods section. The ROIs were delineated to dissect streamlines that terminate in the region surrounding the PPA on one end, and in the medial occipital lobe (cuneus and lingual gyrus) on the other end. These dissections confirmed the existence of a large, coherent white matter bundle connecting EVC in the medial occipital lobe to the posterior MTL. On one end, most of the streamlines of this bundle project in the anterior portions of areas V1, V2 and V3 within the occipital lobe. On the other end, most streamlines terminate within the posterior portion of the PPA.

Figure 5 shows the 3D reconstruction of this bundle, within the brain surface of an example HCP participant. After leaving the anterior peri-calcarine cortex with a lateral course, fibres emerging from the cuneus (Cu) and lingual gyrus (LG) merge into a single bundle. This bundle continues anteriorly toward the temporal lobe and terminates in the region that overlaps the anterior tip of the LG and the posterior PHG. The entire course of this bundle is infero-medial to the occipital horn and atrium of the lateral ventricles. We chose to label this bundle as the medial occipital longitudinal tract (MOLT) owing to its location and anatomical course.

**Figure 5.**
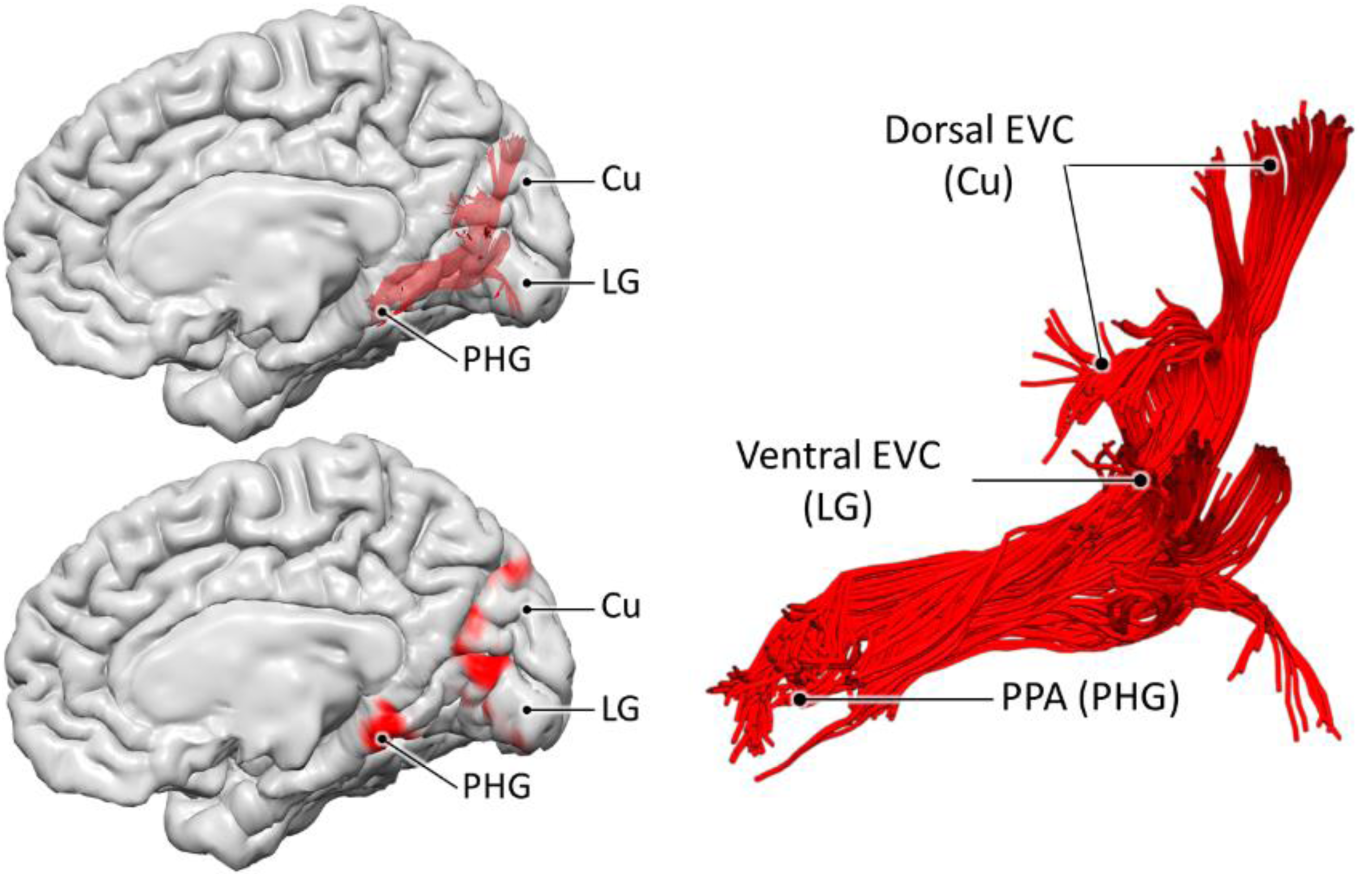
The medial occipital longitudinal tract (MOLT). Tractography reconstruction of the medial occipital longitudinal tract (MOLT) in an example HCP participant. The MOLT is an occipito-temporal white matter pathway that stems from the anterior cuneus (Cu) and lingual gyrus (LG) and terminates in the posterior parahippocampal gyrus (PHG). In the medial occipital lobe, it projects onto peripheral visual field representations within early visual cortex (EVC), while its temporal lobe terminations overlap the posterior parahippocampal place area (PPA).

The distribution of the MOLT’s projections within the occipital lobe suggests the existence of two components within this bundle, and their further anatomical characterisation may provide insight into the MOLT’s functional role. For this reason, we used three tractography-based metrics to assess the macrostructure and microstructure of the Cu and LG components of the MOLT and their interhemispheric lateralisation. The first metric was tract volume measured as the total volume of voxels intersected by streamlines (i.e., spatial occupancy) for each of the Cu and LG components in the two hemispheres. The second metric was the surface area of their cortical projections within EVC. The third metric was the hindrance modulated orientational anisotropy (HMOA), which is a proxy for the microstructural measure of fibre density ^45^. For each metric we calculated a (Cu-LG)/(Cu+LG) ratio within each hemisphere, and a (RH-LH)/(RH+LH) ratio for each component (Figure 6). Full descriptive statistics of these metrics and comparisons are reported in the Supplemental Material, and more details about these comparisons are available in the Methods section.

**Figure 6.**
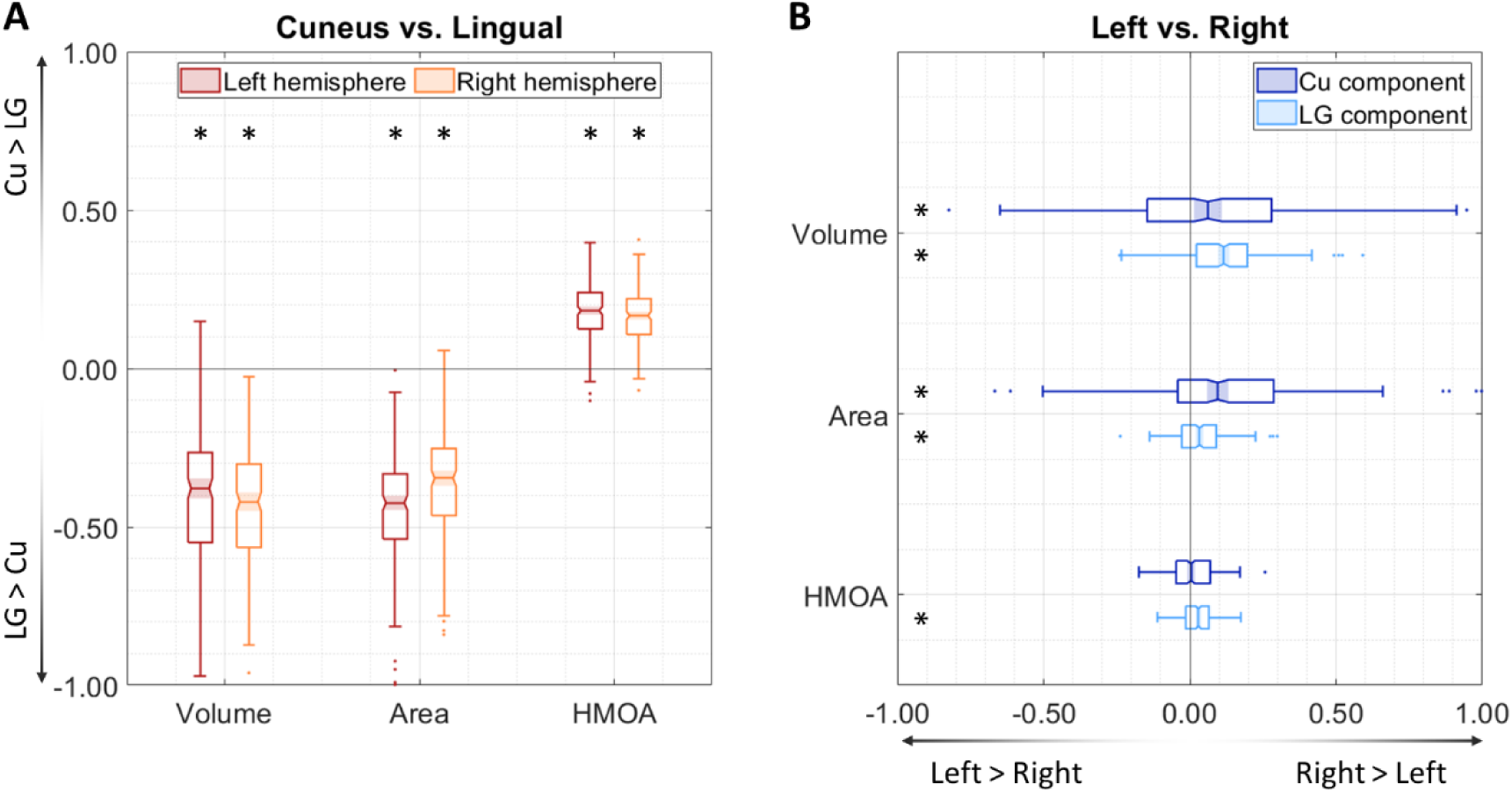
Macrostructural and microstructural assessment of the MOLT. These box charts summarise the comparisons between the MOLT’s components within each hemisphere and across hemispheres in 200 subjects. **(A)** In both hemispheres, the lingual gyrus (LG) component has a larger volume, distributes to a wider occipital surface, and has a lower fibre density (as indicated by its lower HMOA) compared to the cuneus (Cu) component. **(B)** Both the Cu and LG components of the MOLT tend to exhibit a slight lateralisation towards the right hemisphere. Within the box charts, the notch and shaded bar represent the median value, and the edges of the box correspond to the upper and lower quartiles. Asterisks indicate that the mean of the corresponding distribution is significantly different from zero after Bonferroni correction (statistics are fully reported in the Supplementary Material).

The comparison between the MOLT components within each hemisphere revealed that the LG component is larger in size and targets a larger surface area within EVC compared to the Cu component. The LG component also has lower HMOA values, which possibly indicates a lower axonal cohesion and/or density compared to the Cu component. The inter-hemispheric comparison revealed that both Cu and LG components of the MOLT are larger in size, project to a larger surface area within EVC, and have higher HMOA values in the right hemisphere. Overall, these results indicate that the MOLT may carry more visual information about the upper visual field and have a higher information capacity in the right hemisphere.

## DISCUSSION

### PPA clusters with different connectivity profiles

Current models of visuo-spatial processing ^21,29^ focus on input to the MTL via multiple routes. Input that is specific to the PPA seems to arise mainly from indirect parietal projections relayed via the RSC/MPC, or from direct projections from area V4 of EVC. This view does not disagree with our findings that different sub-zones within the PPA receive information via separate anatomical pathways. On one hand, we observe that the anterior zone (aPPA) is strongly connected to the RSC/MPC, which is a connectivity profile that closely agrees with the descriptions of the parieto-medial-temporal branch of the dual-stream model ^21^ and previous functional connectivity analyses of this pathway ^46,47^. On the other hand, the posterior zone (pPPA) is strongly connected to the anterior-most portions of the Cu and LG, overlapping peripheral visual field representations within EVC. This observation challenges the current view that EVC projections to the PPA mainly arise from V4, and instead suggests that earlier visual areas contribute directly to PPA afferents.

The idea that the PPA contains multiple subunits with different connectivity profiles has already been alluded to in the literature. Cytoarchitectonic data suggests that TFO, the posterior-most area of the macaque PHG, has a prominent layer IV and a general laminar profile more akin to the profiles of visual areas than to those of its anterior neighbours TF and TH ^48^. Further, evidence from task fMRI studies of the human brain suggests that, within the PPA, the posterior portion exhibits stronger functional coupling with peripheral representations within EVC, while the anterior portion shows a strong coupling with an extended fronto-parietal network ^17,18^. Additionally, the pPPA contains two distinct retinotopic maps of the visual field, PHC1 and PHC2 ^49^, which do not extend to its anterior-most border defined by anatomo-functional approaches ^39^.

This view of the PPA might seem different from its original description as a cortical region which shows greater functional responses to images of scenes compared to those of isolated faces or objects ^12^. Yet, the two views go together given that the main features in a visual scene are its space-defining borders and the spatial configuration of its constituent elements ^16,17,50^. Additionally, the PHG plays a general role in processing contextual associations, with spatial associations activating the pPHG and non-spatial associations activating the anterior PHG ^10,51,52^.

### The medial occipital longitudinal tract (MOLT)

In our analysis of 200 HCP subjects, we consistently replicated a coherent white matter bundle that runs between the peripheral visual representations within EVC and the PPA. Considering its medial location and course along the posterior-anterior axis, we refer to this pathway as the medial occipital longitudinal tract (MOLT). The MOLT has a dorsal (Cu) and a ventral (LG) component that both project onto the same zone in the posterior PPA. Fibres of the MOLT were identified in early tractography studies but considered as part of the ventral cingulum ^53^ and inferior longitudinal fasciculus ^54^. The existence of a direct connection between EVC and posterior PHG as a separate tract from the cingulum was presented by Catani ^33^ and indicated with the descriptive term ‘Sledge Runner’ fasciculus. Recent *post mortem* dissection and *in vivo* tractography studies confirmed the existence of the ‘Sledge Runner’ fasciculus but they limited its occipital terminations to the most dorsal portion of the cuneus ^34,35,55^. Therefore, we believe that the ‘Sledge Runner’ fasciculus should only be considered as the dorsal-most component of the MOLT, possibly projecting onto dorsal V2 and V3.

This distinction becomes more important when we consider that the MOLT mediates a stronger overall connectivity between the PPA and ventral EVC (LG, upper visual field) compared with dorsal EVC (Cu, lower visual field). This imbalance is in line with a functional bias within the PPA which contains a larger representation of the upper visual field ^29,49,56^. The MOLT also exhibits a right hemisphere lateralisation, which is an observation that fits existing literature reporting such a hemispheric bias in the spatial learning functions supported by the parietal and medial temporal lobes ^36,57^. Further, this lateralisation may explain the higher frequency of visuospatial learning deficits following posterior right hemisphere lesions ^6,57^.

Axonal tracing reports in the macaque using anterograde tracer injections in EVC, specifically in peripheral visual field representations in V2, lead to traces in the posterior PHG ^31^. Other reports using anterograde injections in the PHG itself show traces in the peri-calcarine occipital region ^32^. This suggests that bidirectional information exchange takes places between the posterior PHG and EVC in the macaque. Therefore, the MOLT likely represents reciprocal anatomical connections between EVC and the PPA in the human brain.

### Role of the MOLT in visuospatial learning

According to clinical reports, lesions affecting the posterior PHG and anterior tip of the LG (coinciding with the posterior PPA cluster) lead to a deficit known as landmark agnosia in which patients are “unable to represent the appearance of salient environmental stimuli” ^57^. Conversely, lesions to more anterior MTL regions and/or RSC lead to anterograde disorientation, meaning that patients are “unable to create new representations of environmental information” ^57^. This suggests that the posterior cortical regions are more important for encoding the local space, while the anterior regions are involved in placing and retrieving this information within the context of existing knowledge. Indeed, through the use of meta-analytic maps associated with these two terms, we found that the posterior PPA is more involved in encoding, while retrieval is only present in the anterior PPA, in line with previous reports ^24,41,43^.

Our view is that the MOLT carries ‘raw’ visual information from EVC which the PPA requires, in combination with higher-order spatial information stemming from the parietal lobe, to fully map the visual scene. In other words, afferents from EVC and from RSC/MPC must work in tandem to allow the PPA to fully carry out its role. In this context, the MOLT may carry feedforward and feedback spatial information between EVC and the PPA, thereby serving as a pathway for re-entrant visual information ^58^ that supports a multistage encoding and learning of the visual scene. As such, early activations within the PPA ^26^ may correspond to an initial mapping of the ‘gist of the scene’ which, via feedback to EVC, refines the later, detailed mapping of object configuration. This refined information, still in a retinotopic space, could then be combined with parietal and retrosplenial spatial information at longer latencies ^27^, and translated into an allocentric frame of reference. This higher order, viewpoint-invariant spatial information would ultimately reach more anterior MTL regions including the hippocampal formation, and more distant frontal regions ^21,28,32,59^.

### Limitations and future steps

Here, we applied a multimodal investigation to a large cohort of healthy participants. It remains important to list some limitations and suggestions for future investigations. First, neuroimaging methods, including fMRI and diffusion tractography, rely on models to produce results. Although these depend on data quality and choice of processing pipeline, we used high resolution data processed according to best practices to minimise such biases. Second, future investigations aiming to ascertain the role of the MOLT in the spatial learning domain would benefit from including specific testing for its putative function along with imaging investigation within the same cohort. Finally, the data we used is from a young adult cohort, so future investigations of the MOLT’s developmental trajectory may offer insights into the development of visuospatial learning abilities across the lifespan.

### Conclusion

Based on converging evidence from structural and functional connectivity, and from the distribution of the encoding and retrieval systems within the PPA, we believe that the MOLT serves as a ventral white matter pathway that carries information crucial for visuospatial learning.

## ARTICLE DETAILS

## Acknowledgements

This work was supported by a Wellcome Trust Investigator Award (No. 103759/Z/14/Z). Data were provided in part by the Human Connectome Project, WU-Minn Consortium (Principal Investigators: David Van Essen and Kamil Ugurbil; 1U54MH091657) funded by the 16 NIH Institutes and Centers that support the NIH Blueprint for Neuroscience Research; and by the McDonnell Center for Systems Neuroscience at Washington University. The authors are grateful to Dr Carlo Sestieri, Dr Massimo Caulo, Dr Giuseppe Zappalà, Dr Sergio Della Sala, and the members of the NatBrainLab (www.natbrainlab.co.uk) for their feedback.

## Author contributions

Conceptualization, A.B., F.D.A, D.f., and M.C.

Methodology, A.B., F.D.A., D.C., F.D.R., D.f., and M.C.

Software development, A.B., F.D.A., D.C., and F.D.R.

Investigation, A.B., F.D.A., and M.C.

Resources, A.B., F.D.A., and M.C.

Writing – Original Draft, A.B. and M.C.

Writing – Review & Editing, A.B., F.D.A., D.C., F.D.R., D.f., and M.C.

Funding Acquisition, M.C.

## METHODS

### MRI Data

DWI data from 200 healthy participants (100 females; age = 29.16 ± 3.73 years) were obtained from the Human Connectome Project (https://www.humanconnectome.org). All subjects were right-handed as determined by a score of 50 and above on the Edinburgh handedness questionnaire ^60^.

Diffusion MRI data of the HCP were acquired on a 3 T Siemens “Connectome Skyra” using a spin-echo EPI sequence (TR = 5520 ms; TE = 89.5 ms; matrix of 168×144; 111 slices with a thickness of 1.25 mm; isotropic voxels of 1.25 mm; multiband factor = 3). Three diffusion-weighted shells were acquired (*b* = 1000, 2000 and 3000 s/mm^2^) with two opposite phase-encoding directions each (L>>R and R>>L). Each shell consisted of 90 diffusion-weighted directions and six interleaved non-diffusion-weighted volumes. For this study, only data from the *b* = 2000 s/mm^2^ shell were used as this b-value offers a good compromise between signal-to-noise ratio and high angular resolution, especially at high spatial resolution ^61^.

The data were obtained from the HCP database in pre-processed form following the HCP minimal pre-processing pipelines ^62^. Briefly, correction for motion and eddy current distortions was performed using *eddy* with outlier slice replacement ^63,64^. Correction of susceptibility distortions was incorporated into this step by means of an off-resonance field estimated using *topup* ^65^. Diffusion MRI data were then corrected for gradient non-linearity and finally aligned to the structural space using a boundary-based registration ^66^.

Tractography was computed using StarTrack (https://www.mr-startrack.com/). Spherical deconvolution was based on a damped version of the Richardson-Lucy algorithm ^45,67^ with the following parameters: fibre response *α* = 1.8; number of iterations = 300; amplitude threshold *η* = 0.0020; geometric regularisation *ν* = 12. Fibre tracking was then performed using the multi-fibre Euler-like algorithm with the following parameters: minimum HMOA threshold = 0.0033; step size = 1.0 mm; maximum angle threshold = 45°; minimum fibre length = 20 mm; maximum fibre length = 300 mm.

### Building vertex-wise connectivity matrices

The whole-brain tractogram of each subject was converted to a vertex-wise structural connectivity matrix. For each subject, this approach was based on the native space midthickness cortical mesh, vertex-matched to the ‘32k FS LR’ surface template using the multimodal surface matching (MSM) method ^38,68^. These surfaces have the advantage of maintaining the native anatomy of the brain while offering a vertex-level matching between subjects. As a result, information from multiple subjects can be directly compared at each vertex.

The end points of each tractography streamline were projected to the nearest vertex on the midthickness surface, with a maximum allowed distance of 4 mm. Only streamlines that resulted in two cortical targets (one for each end point) survived this step. Due to the sparse distribution of streamlines near the cortical surface, this approach would result in a patchy representation of cortical targets which is problematic for group-level analysis. To compensate for this and for tractography’s uncertainty near grey matter ^69^, a geodesic Gaussian kernel (FWHM = 2.5 mm) was applied to each target independently. The connectivity information resulting from each streamline was used to populate a 32492×32492 sparse matrix representing all vertices of the cortical surface. Each subject’s matrix was then normalised by its highest value before the group-level mean connectivity matrix was computed.

### Defining cortical ROIs

In preparation for the connectivity analyses in the following sections, several regions of interest were defined using published cortical atlases.

First, an ROI was defined to cover EVC and included the following labels from the multimodal parcellation (MMP) atlas ^38^: V1, V2, V3, V4, V3A, V6. This effectively covered the Cu, LG and occipital pole, and included all eccentricity representations within EVC. Although the V3A and V6 regions are not EVC per se, they were included in this ROI to ensure the ROI’s spatial continuity and avoid introducing sharp boundaries that may affect the clustering analysis.

A second ROI was defined to cover the MPC, RSC, and cortical areas that relay information from the parietal lobe to the MTL according to the dual-stream model ^21^ and are important for spatial learning based on human case studies ^36^. It included the following MMP atlas labels: ProS, DVT, POS1, POS2, 7m, 7Pm, 7Am, PCV, 31pd, 31pv, 31a, d23ab, v23ab and RSC. Similar to the EVC ROI, this ROI also included other neighbouring labels of the MMP atlas to ensure its spatial continuity and that no artefactual boundaries are created.

A third ROI was defined to cover the entire length of the MTL and included the following MMP atlas labels: VMV1, VMV2, VMV3, PHA1, PHA2, PHA3, VVC, EC, PreS, H and PeEc. These labels mainly span the length of temporal cortex medial to the collateral sulcus.

Finally, an ROI covering the PPA was obtained from Weiner et al. ^39^ who demonstrated that the location of functional responses within the PPA can be predicted with high accuracy using surface topology alone. The ROI covered parts of the posterior CoS and PHG, and anterior LG. This ROI partially overlaps the following labels from the MMP atlas ^38^: VMV1, VMV2, VMV3, PHA1, PHA2 and PHA3. The ROI was released by the authors in the form of a FreeSurfer label registered to the ‘fsaverage’ template, which we subsequently realigned to the ‘32k_FS_LR’ template following HCP guidelines.

### Connection density in the MTL

The aim of this analysis was to assess the spatial distribution of the density of connections stemming from the EVC and RSC/MPC ROIs, and projecting within the MTL. To this end, the vertex-wise connectivity matrices previously computed were filtered to only keep connections between the MTL and either of these ROIs. The obtained values for the MTL vertices were then sorted according to the position of each vertex along the y-axis and grouped into 1 mm bins. The connectivity value was first computed for each subject separately in each bin, then the group mean and standard deviation were calculated. This effectively allowed for the assessment of the relationship between the location of an MTL vertex along the anterior-posterior axis and the strength of its connectivity to the EVC or RSC/MPC ROIs. A Spearman rank correlation was finally computed to assess the relationship between projection density and position along the y-axis.

### PPA clustering and connectivity analysis

The aim of this analysis was to assess whether the PPA contains multiple sub-units with different anatomical connectivity profiles based on a data-driven approach. First, the values of the subject-level connectivity matrices previously computed were converted to z-scores ^70^, and the mean z-matrix was computed. This matrix was then filtered to only retain the connections between the PPA on one end, and both the EVC and RSC/MPC ROIs on the other end.

The mean z-matrix was entered into a principal component analysis (PCA). In this context, the z-matrix acted as a large dataset where the PPA vertices represented variables and the EVC/RSC/MPC vertices represented observations of these variables. This approach was blind to the locations of vertices on the brain surface and was thus completely driven by the connectivity profile of each PPA vertex. A scree plot ^71^ was used to determine the number of components to retain. The resulting principal components effectively represented the PPA region in a reduced space of only a few variables. Importantly, this step also acted as a filter that suppressed contributions from ‘noisy’ connections, or ones which had very high inter-subject variability.

Agglomerative hierarchical clustering (Euclidean distance, ward algorithm) was applied to the resulting principal component coefficients to extract binary clusters within the PPA. This approach resulted in a dendrogram (hierarchy tree) where each observation (PPA vertex) was linked to other observations in a hierarchical fashion. These links defined clusters of vertices that were most similar in their loadings on the principal components. The spread vs. separation (SS) index proposed by Moreno-Dominguez et al. ^40^ was used to objectively determine the optimal number of clusters. A higher SS index means that the variability of observations between clusters is greater than the variability within clusters. Therefore, the number of clusters was chosen to correspond to the highest SS index.

The mean structural connectivity of each resulting cluster was then obtained from the mean connectivity matrix previously computed. Additionally, the mean functional connectivity of each cluster was calculated based on the group average dense resting-state functional connectome released by the HCP as part of the ‘HCP-S1200_GroupAvg_v1’ dataset (https://www.humanconnectome.org/study/hcp-young-adult/document/extensively-processed-fmri-data-documentation). Tract-based connectivity values, which follow a heavily skewed distribution compared to functional connectivity, were log-transformed ^40,72^. Spearman rank correlations (one-tailed) were then computed between tract-based and functional connectivity measures for each cluster to assess the degree of agreement between the two modalities.

### Functional meta-analysis

Two association test maps were generated for the terms ‘encoding’ and ‘retrieval’ following a meta-analytic approach ^42^ using NeuroSynth (https://neurosynth.org/). For each voxel, these maps contain a z-score from a two-way ANOVA indicating how consistently that voxel is activated in studies that mention the key term compared with studies that do not. These maps are corrected for multiple comparisons using a false discovery rate of .01. Each map was projected to the surface of the MNI brain, allowing us to use the PPA mask as an ROI for the analysis. The maximum z-score for each coordinate along the PPA’s anterior-posterior axis was plotted for each of the two maps, allowing us to compare their distribution along the PPA.

### Virtual dissections

Virtual dissections were performed in TrackVis (http://trackvis.org/) to extract the anatomical bundle that gave rise to the observed connectivity patterns between the posterior PPA cluster and the EVC in the clustering analysis.

MegaTrack ^44^, a supervised semi-automatic group-level approach, was used to handle the large number of participants. Each participant’s anisotropic power map ^73^ was mapped to the space of the MNI152 template using the symmetric normalisation algorithm in ANTs ^74,75^. The resulting affine and diffeomorphic warping were applied to each participant’s tractogram and all tractograms were concatenated to create a single ‘mega’ tractography dataset. After performing virtual dissections of the large dataset in common space, each participant’s streamlines were mapped back to native space where tract-specific measurements could be extracted, and cortical projections analysed.

To isolate the tracts of interest, an ROI was manually defined on sagittal and coronal slices to intersect any streamlines entering the medial occipital lobe, covering the LG and Cu. A sphere ROI was defined at the intersection between the LG and PHG, overlapping the anatomical location of the PPA. Only streamlines terminating within the PPA ROI and crossing the occipital ROI were retained. Additional ROIs were used to manually refine the dissected tracts and remove spurious streamlines (e.g., streamlines that took unusually long trajectories or crossed sulcal boundaries).

For further analysis of the subcomponents of the dissected bundle, two additional occipital ROIs were defined to isolate the component projecting to the cuneus and the one projecting to the lingual gyrus. To this end, the cuneus ROI was defined to include any streamlines of the dissected bundle which terminate above the calcarine sulcus, and similarly the lingual ROI was defined to include streamlines terminating below the calcarine sulcus.

### Assessment of lateralisation and vertical bias in the dissected tracts

Hemispheric lateralisation for each of the dorsal and ventral components was assessed according to the following formula:

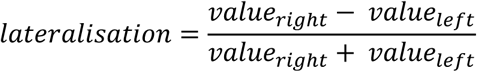

where *value* represents one of the following metrics: (1) tract volume; (2) surface area of connected EVC cortex; (3) hindrance modulate orientational anisotropy (HMOA). In this way, a lateralisation toward the right hemisphere would lead to positive values, and left lateralisation would lead to negative values.

Additionally, given that that PPA contains a substantially larger representation of the upper visual field ^49,56^, the connectivity between the PPA and ventral EVC (LG, representing the upper visual field) was expected to be stronger than that with dorsal EVC (Cu, representing the lower visual field). This vertical bias was assessed following a similar approach to the one used for lateralisation, according to the following formula:

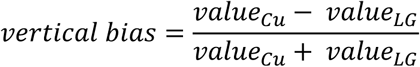

where *value* also represents one the previously described metrics, and where a dominance of the Cu component would result in positive values and a dominance of the LG component would result in negative values.

For each of these comparisons, statistical significance was determined through a one sample t-test (two-tailed) performed on the resulting bias index. The Bonferroni corrected significance level was set to 0.0042.

