## Supplemental Material for "The missing pathway in current visuospatial processing models"

**Further analysis of the connectivity profiles of the three PPA clusters**

In our main analysis, we found overall good agreement between the structural and functional connectivity profiles of the posterior and anterior parahippocampal place area (PPA) clusters, but the results were more ambiguous for the lateral cluster. Therefore, we performed further analysis to generate intramodal comparisons for each pair of clusters. To this end, for each pair of clusters, we ran a Spearman rank correlation based on the connectivity values of those two clusters to each vertex of the large ROI encompassing early visual cortex (EVC), retrosplenial cortex (RSC), and medial parietal cortex (MPC). The results of these comparisons are summarised in Figure S1 for structural connectivity, and in Figure S2 for functional connectivity.

Note that because these comparisons are intramodal, weak or moderate correlations bear no substantial value. This is more so the case for functional connectivity due to its lower spatial specificity. This means that the functional connectivity profiles of two clusters should be considered similar only if they are strongly correlated.

The results of the structural connectivity comparisons show no correlations or weak correlations between the observed connectivity profiles of the different PPA clusters (Figure S1). Naturally, this should be expected given that the initial hierarchical clustering was based on this information, rendering this a circular question. Comparisons of the functional connectivity values of pairs of clusters corroborate these observations for the posterior and anterior PPA clusters (Figure S2). However, the posterior and lateral clusters were indeed strongly correlated in both hemispheres.

To further understand the similarities and differences between the three clusters in an anatomical context, we subtracted the connectivity values of each pair of clusters (within each modality) and plotted the resulting maps on the brain surface (Figure S3). As expected, the structural and functional difference maps lend further support to the division between the anterior and posterior PPA clusters: the posterior cluster had a clear preference for EVC, while the anterior cluster showed a clear preference for the RSC/MPC. The comparison of the posterior and lateral clusters revealed that the posterior PPA's structural connectivity to anterior EVC is stronger than that of the lateral PPA, in both hemispheres. Further, although there is almost no difference between the two clusters based on functional connectivity in the left hemisphere, the posterior cluster is more functionally connected to anterior EVC than the lateral cluster in the right hemisphere (albeit the difference is not very large).

To explain the observed discrepancy between the two modalities, we need to turn to the anatomical properties of the region surrounding the PPA. As functionally defined, the PPA sits in the collateral sulcus, with one portion spanning its medial wall, i.e., the parahippocampal gyrus, and one portion spanning its lateral wall, i.e., the fusiform gyrus. This raises two possible scenarios that may drive the observed functional and structural connectivity profiles. The first scenario relates to the limited ability of fMRI to distinguish the BOLD signal originating from the two walls of a sulcus. fMRI suffers from partial volume effects and mixing signal from neighbouring gyri. It is therefore possible that the PPA's functional localisation encroaches on the lateral wall of the Cos more than the real underlying neuronal populations do. The second possible scenario is that tractography is more capable of visualising connections between EVC and the medial wall of the collateral sulcus. This is especially possible given the known limitations of tractography in reaching deep sulcal locations, an effect known as the 'gyral bias' [1]. In this case, the collateral sulcus may form an artificial boundary for tractography which drives the clustering to show two different zones in the posterior side of the PPA.

It is difficult to determine which of these two scenarios is driving the observed difference between structural and functional connectivity for the lateral PPA cluster, and it is likely that the data is affected by both to some extent. However, the fact that the posterior cluster shows more functionally connected to the anterior EVC in the right hemisphere only (Figure S3, second column) indicates that there is indeed a functional difference between the two clusters and that they are not identical.

1. Schilling, K. *et al.* Confirmation of a gyral bias in diffusion MRI fiber tractography. *Hum. Brain Mapp.* **39**, 1449–1466 (2018).

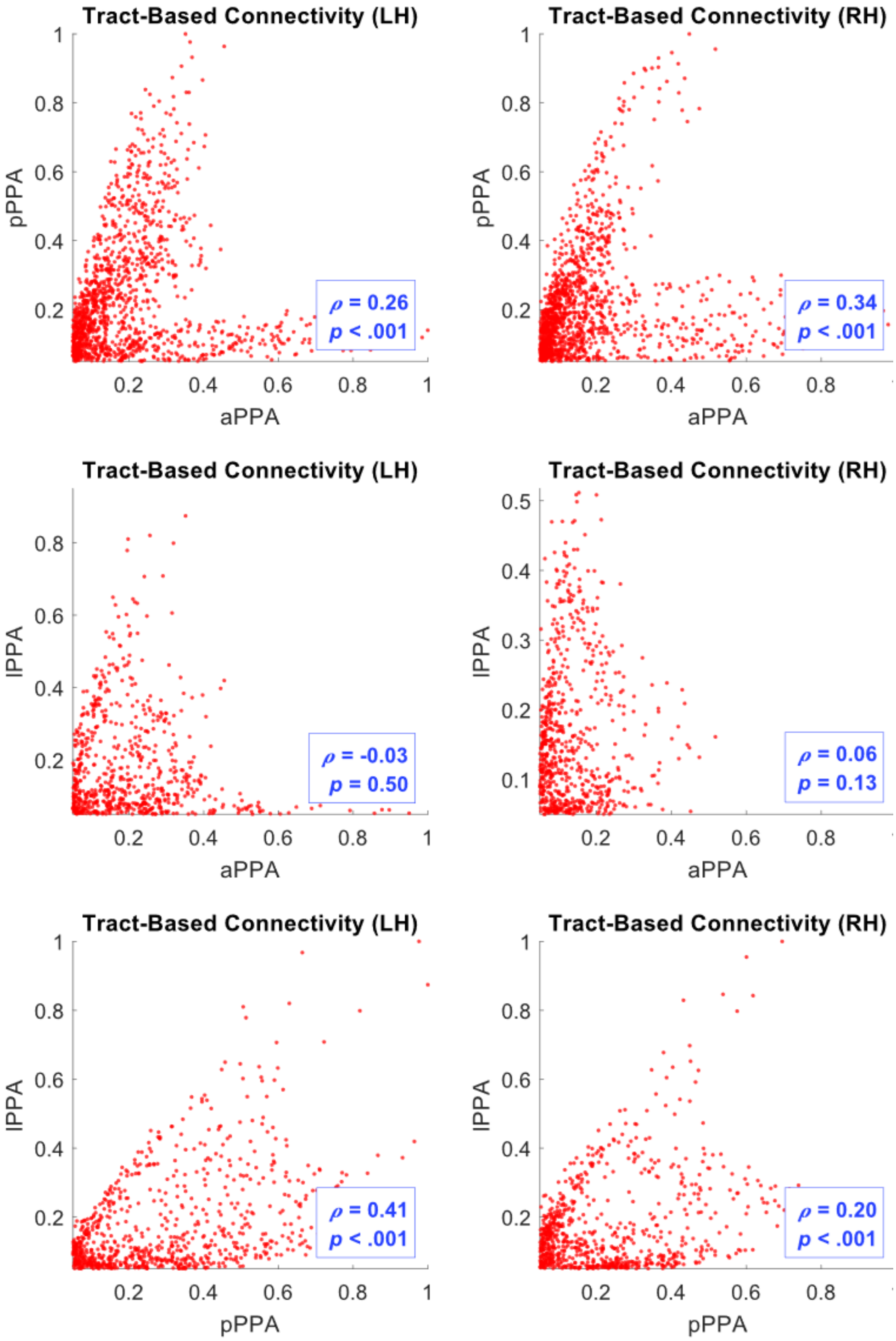


**Figure S1. Intramodal comparison of the structural connectivity of PPA clusters.**

Each chart plots tract-based connectivity values for one PPA cluster against those of another. The text boxes show Spearman rank correlations.

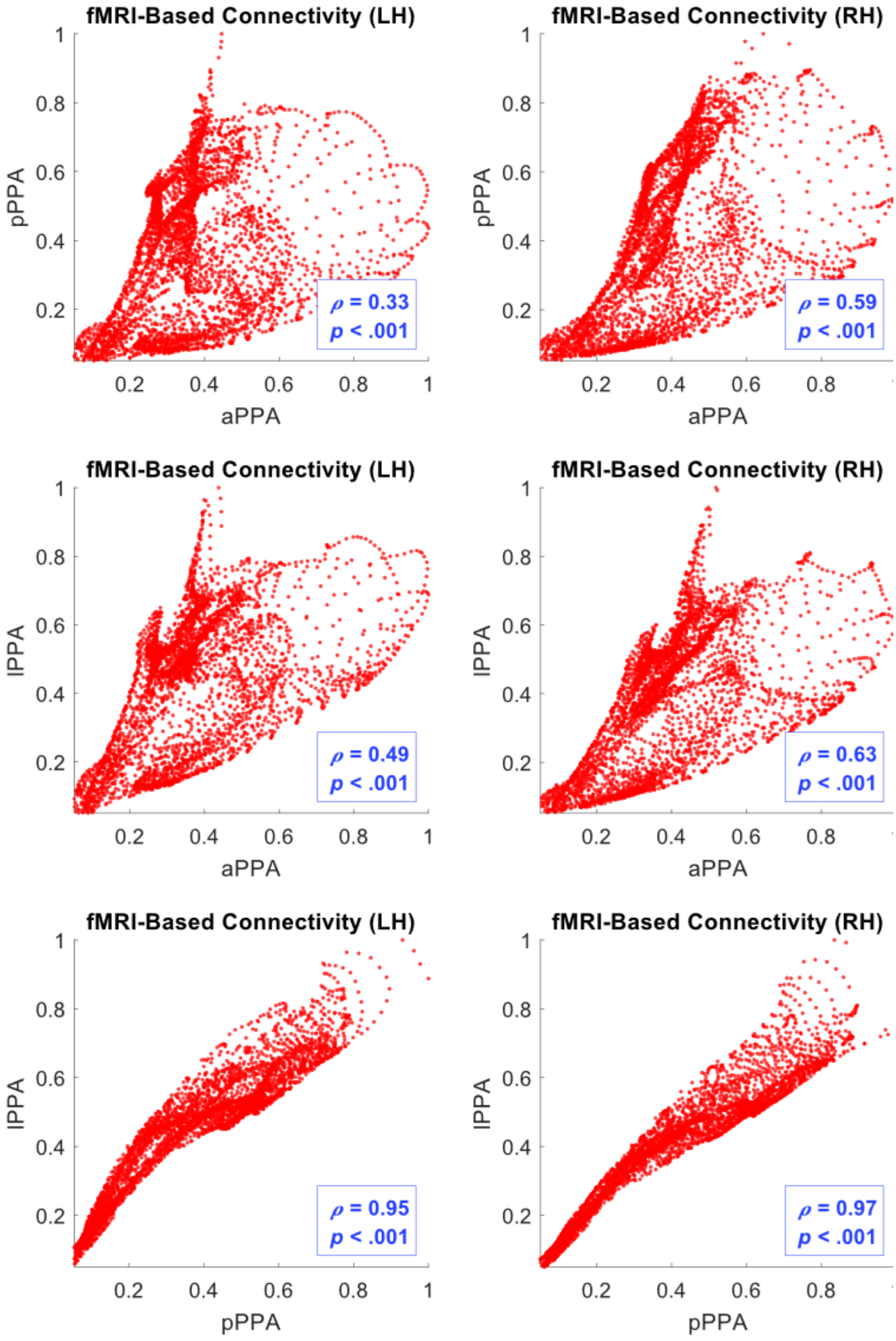


**Figure S2. Intramodal comparison of the functional connectivity of PPA clusters.**

Each chart plots fMRI-based connectivity values for one PPA cluster against those of another. The text boxes show Spearman rank correlations.

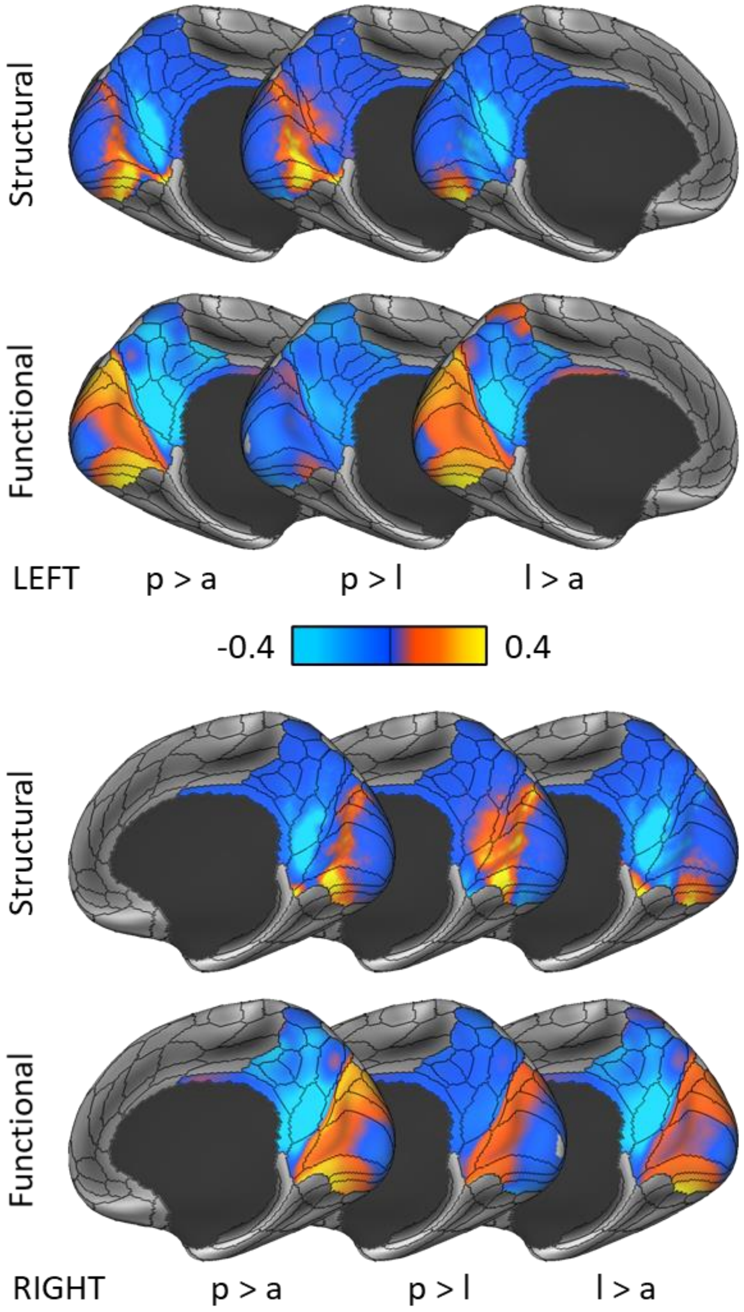


**Figure S3. Intramodal comparison of the connectivity profiles of each pair of PPA clusters on the brain surface.**

To further clarify the similarities and differences between the connectivity profiles of the three PPA clusters, each pair of clusters were additionally compared by direct subtraction. Despite the apparent functional similarity between the posterior and lateral clusters previously found, the posterior cluster still shows higher functional connectivity to anterior EVC in the right hemisphere compared with the lateral cluster. p: pPPA; l: lPPA; a: aPPA.

**Table S1. Descriptive statistics of the medial occipital longitudinal tract (MOLT).**

| | *Streamline count* | *Tract volume (millilitres)* | *Connected area (mm2)* | *HMOA* |
|---|---|---|---|---|
| *MOLT Dorsal Left* | 126.84 ± 111.24 | 3.33 ± 1.76 | 1360.19 ± 544.86 | 0.0147 ± 0.0026 |
| *MOLT Ventral Left* | 265.28 ± 117.33 | 7.45 ± 2.20 | 2817.19 ± 546.70 | 0.0101 ± 0.0010 |
| *MOLT Dorsal Right* | 169.84 ± 125.70 | 3.74 ± 1.65 | 1681.94 ± 560.25 | 0.0149 ± 0.0024 |
| *MOLT Ventral Right* | 368.38 ± 176.45 | 9.24 ± 2.43 | 3048.01 ± 522.36 | 0.0106 ± 0.0011 |

**Table S2. Statistical comparison of the left and right hemisphere MOLT components.***

| | *Mean* | *95% CI* | *t* | *df* | *p* |
|---|---|---|---|---|---|
| *MOLT Cu HMOA* | 0.01 | -0.01, 0.02 | 1.27 | 198 | .205 |
| *MOLT Cu Volume* | 0.07 | 0.01, 0.14 | 3.15 | 198 | .002 |
| *MOLT Cu Area* | 0.11 | 0.06, 0.17 | 6.20 | 198 | < .001 |
| *MOLT LG HMOA* | 0.02 | 0.01, 0.03 | 6.03 | 199 | < .001 |
| *MOLT LG Volume* | 0.11 | 0.08, 0.14 | 10.98 | 199 | < .001 |
| *MOLT LG Area* | 0.04 | 0.02, 0.06 | 6.68 | 199 | < .001 |

** Positive values indicate a right hemisphere lateralisation.*

**Table S3. Statistical comparison of the dorsal (cuneus, Cu) and ventral (lingual gyrus, LG) MOLT components.***

| | *Mean* | *95% CI* | *t* | *df* | *p* |
|---|---|---|---|---|---|
| *Cu vs LG Left HMOA* | 0.18 | 0.16, 0.20 | 29.69 | 198 | < .001 |
| *Cu vs LG Left Volume* | -0.41 | -0.45, -0.36 | -26.66 | 198 | < .001 |
| *Cu vs LG Left Area* | -0.37 | -0.40, -0.33 | -29.70 | 198 | < .001 |
| *Cu vs LG Right HMOA* | 0.16 | -0.45, -0.31 | -15.70 | 199 | < .001 |
| *Cu vs LG Right Volume* | -0.43 | -0.47, -0.39 | -32.12 | 199 | < .001 |
| *Cu vs LG Right Area* | -0.30 | 0.15, 0.18 | 28.67 | 199 | < .001 |

** Negative values indicate a dominance of the LG component.*
